# Africanized honey bees in Colombia exhibit high prevalence but low level of infestation of *Varroa* mites and low prevalence of pathogenic viruses

**DOI:** 10.1101/2020.12.21.423754

**Authors:** Víctor Manuel Tibatá, Andrés Sanchez, Evan Young-Palmer, Howard Junca, Victor Manuel Solarte, Shayne Madella, Fernando Ariza, Judith Figueroa, Miguel Corona

## Abstract

The global spread of the ectoparasitic mite *Varroa destructor* has promoted the spread and virulence of highly infectious honey bee viruses. This phenomenon is considered the leading cause for the increased number of colony losses experienced by the mite-susceptible European honey bee populations in the Northern Hemisphere. Most of the honey bee populations in Central and South America are Africanized honey bees, which are considered more resistant to *Varroa* compared to European honey bees. However, the relationship between *Varroa* levels and spread of honey bee viruses in Africanized honey bees remains unknown. In this study, we determined *Varroa* prevalence and infestation levels as well as the prevalence of seven major honey bee viruses in Africanized honey bees from three regions of Colombia. We found that although *Varroa* exhibited high prevalence (92%), its infestation levels were low (4.6%) considering that these populations never received acaricide treatments. We also detected four viruses in the three regions analyzed, but all hives were asymptomatic, and virus prevalence was considerably lower than those found in other countries with higher rates of mite-associated colony loss (DWV 19.88%, BQCV 17.39%, SBV 23.4 %, ABPV 10.56%). Our findings indicate that AHBs possess natural resistance to *Varroa* that does not prevent the spread of this parasite among their population, but restrains mite population growth and suppresses the prevalence and pathogenicity of mite-associated viruses.

## Introduction

Colombian beekeepers have primarily used Africanized honey bees (AHBs) derived from *A. mellifera scutellata* since they arrived in Colombia in 1979 [1, 2] after they escaped from Brazil in 1956 [3]. However, many local beekeepers abandoned this practice after the arrival of AHBs, mainly due to their higher defensiveness. Subsequent generations of Colombian beekeepers adapted their management techniques to deal with the defensiveness of AHBs and at the same time, take advantage of their positive characteristics, including noticeably increased resistance to *Varroa* infestation [4, 5]. This ectoparasite arrived in Colombia in the 1980s, and spread throughout all continental territories, excluding the isolated San Andrés islands located in the Atlantic Ocean.

Honey bee populations in the Northern Hemisphere have experienced severe losses in recent years [6]. These losses have been primarily attributed to infestations by the ectoparasitic mite *V. destructor* [7, 8]. In addition to the direct harmful effects on honey bee health, *V. destructor* is an effective vector for pathogenic honey bee viruses [9–11] Accumulative evidence indicates that the association between *V. destructor* and honey bee viruses, especially Deformed wing virus (DWV), play a crucial role in colony losses [7, 8, 10, 12, 13]. Currently, twenty-three known viral pathogens affect honey bees in different parts of the world [14]. The seven most widely distributed and pathogenic viruses include DWV, sacbrood virus (SBV), black queen cell virus (BQCV), acute bee paralysis virus (ABPV), chronic bee paralysis virus (CBPV), Israeli acute paralysis virus (IAPV) and Kashmir bee virus (KBV) [15–18]. In Latin America, published reports on the detection of DWV, CBPV, ABPV, IAPV and SBV exist for Uruguay, Brazil, Argentina, Chile and Mexico [19–28].

Low levels of virus are usually present in honey bees not infected with *Varroa* and persist through horizontal and vertical transmission routes as covert infections [9, 29–32]. However, in apiaries infested with this parasite, especially when *V. destructor* population levels rise, the transmission of viral infections increases, generating overt infections and disease [10, 33–35]. At present, beekeepers in the Northern Hemisphere—who maintain European-derived populations of honeybees—rely on miticides to control *Varroa* infestations and their associated viruses. In contrast, mite control treatments are seldom necessary in South American countries with predominant AHB populations ([21, 25, 36–38] as well as in African countries with populations of *Apis mellifera scutellata* [39]. Results of studies obtained in specific geographic regions of South America and Mexico are consistent with the view that AHBs are more resistant to *Varroa* infestations compared with European honey bees (EHBs) [5, 37, 40–42]. However, this mite still affects AHB colony fitness as reflected by reduced production of honey [42, 43].

The AHB populations in Colombia offers the opportunity to study a natural selection process where populations that are adapted to a tropical climate reproduce and thrive without treatments to control pathogens [44]. The relative resistance of Africanized Honey Bees to *Varroa* infestation might be expected to translate into lower levels of viral infection. However, the prevalence of viruses and their relationship to *Varroa* in Africanized Honey Bees remains inconclusive. For example, one study found no differences in viral prevalence between AHBs and EHBs [38], but another reported increased viral resistance in AHBs compared with EHBs [45]. These conflicting results highlight the need for large-scale field studies to elucidate the epidemiology of *V. destructor* prevalence and infestation levels and their relationship to viral infection in AHB populations, such as those found in Latin American countries [46]. In this study, we determined *V. destructor* infestation levels and the prevalence of seven major honey bee viruses in three regions of Colombia, using honey bee populations that we previously confirmed were composed exclusively of AHBs [44]. Our results enhance knowledge about the relationship between *V. destructor* and viruses in AHBs, and provide the first large-scale field survey of honey bee parasites and pathogens in this altitudinally and seasonally varied equatorial country.

## Materials and methods

### Study design and sampling regions

This study was conducted in three geographical regions located in three representative beekeeping regions of Colombia: Magdalena, Sucre and Boyacá (Figure 1). The number of apiaries, colonies, and municipalities sampled were as it follows: In Magdalena, 151 colonies belonging to eleven apiaries located in six municipalities. In Sucre, 168 colonies from eleven apiaries located in nine municipalities. In Boyacá, 164 colonies from fifteen apiaries distributed throughout twelve counties. The apiaries resided in Tropical and Neotropical regions of Colombia which can have two distinct seasons per year. The “dry season” is characterized by moderate drought, whereas the “rainy season” accounts for most of the annual precipitation. Collections were completed in the Magdalena region during the rainy season (November 2013 and June 2014) and the Sucre region during the dry season (March 2013 and March 2014). Sampling in the Boyacá region deserves special considerations: In the North sub-region, pluvial precipitations present a bimodal tetra-seasonal pattern. In the South, pluvial precipitations exhibit a unimodal bi-seasonal pattern. Thus, timing of the rainy season varies across sub-regions. Samples in Boyacá were collected on the following dates, municipalities and corresponding weather season: October 2013, Soata, Turquemene, Umbita and Guacheta (rainy season); July 2014, San Mateo, Guacheta, Belem, Tutauza (dry season); July 2014, Rondon and Viracacha (rainy season). The study regions encompassed a wide range of altitudes, from near sea level to >3000 m, and ranged in climate from tropical at low altitude to temperate at high altitude. The apiaries in Sucre ranged from an altitude of 26-417 m with a mean of 205 m above the sea level and a tropical climate. Apiaries in Magdalena ranged from 742-1468 m, with a mean of 1053 m and an intermediate, subtropical climate. Apiaries in Boyacá ranged from 2243-3245 m (± 1002 m), <u>with a mean of 2820 m</u> and a temperature climate (supplementary data set 1).

**Figure 1.**
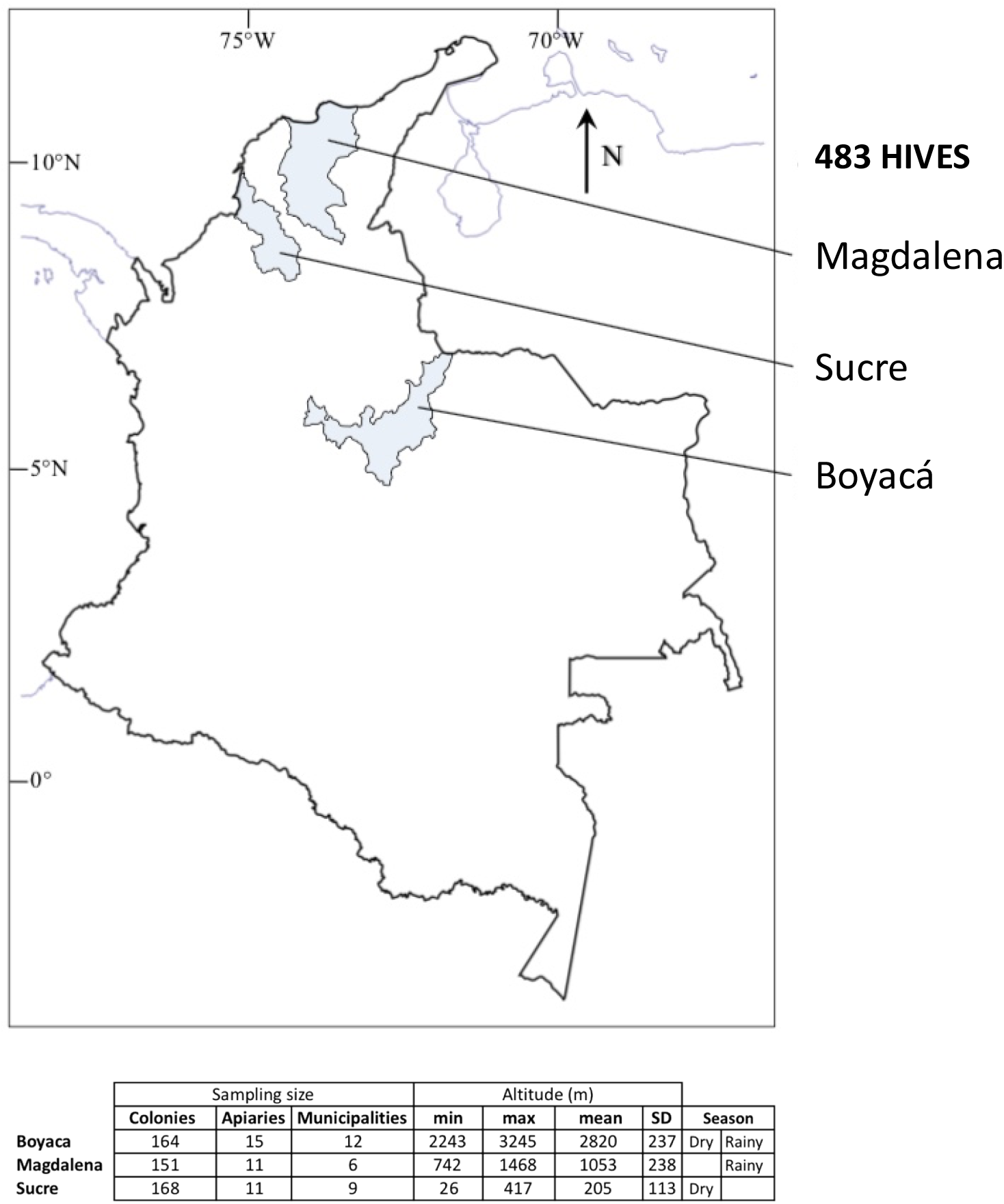
Map of Colombia showing (highlighted in blue) the geographic localization of the three regions analyzed in this study and the number of colonies sampled in each region. Minimum (min), maximum (max), mean and standard deviation (SD) are indicated in meters (m).

Across 5400 colonies from the three regions, 483 were randomly sampled. Sample size estimation was obtained by the program WinEpi from the University of Zaragoza (http://www.winepi.net/) using the formula for known population size (5400 hives), with a confidence level of 97%, considering an expected minimum prevalence of 2% for each of each viral disease. Due to an absence of information about bee morbidity or mortality in the sampled regions, collections were performed randomly. In the sampled apiaries, no symptoms related to viral diseases were observed, and samples where viruses were detected by RT-PCR were considered infected but not necessarily diseased.

### Determination of *Varroa* prevalence and infestation level

In order to determine prevalence and infestation level of *V. destructor* in the 483 hives, 200 adult bees from each colony were deposited in flasks containing 96% ethanol and then transported to the laboratory to obtain the *Varroa* infestation level (VIL) according to the De Jong method [47]. The VIL was calculated as a percentage by dividing the number of *Varroa* mites by the number of bees and multiplied by 100.

### Collection of samples for analysis of RNA viruses

In each of the 483 selected hives, two types of samples were taken: One consisted of 60 larvae and another with 60 adult bees (both taken in pools). The number of larvae and adults per hive was determined by the same formula for the detection of disease. The two samples were collected from a single frame of each hive. Larvae were taken directly from each hive and deposited in 50 mL conical tubes, which contained RNAlater (Qiagen) to avoid RNA degradation. Adult bees were collected alive inside the same hive and were euthanized by inhalation with ethyl acetate in a lethal chamber, according to international standards. All samples were stored and kept in liquid nitrogen until subsequent processing.

### RNA extraction and cDNA synthesis

#### Adults

Each pooled sample of 60 frozen adult bees was placed inside Ziploc bags, and 30 ml of lysis buffer were added (Guanidium thiocyanate 0.8M; ammonium thiocyanate 0,4M; sodium acetate 3M, glycerol 5% and Triton-X 100 2%). The material was smashed with a rolling pin. An aliquot of 620 μl of macerated tissue was taken from each pool, mixed with 320 μl of acid phenol (pH 4.0), incubated 10 min at 95 °C and cooled in an ice bath. Then, 200 μl of chloroform were added. Subsequently, samples were vortexed and centrifuged at 12000 x g for 15 min. at 4°C. The aqueous phase was extracted, and one volume of cold isopropanol was added, mixed and centrifuged at 12000xg for 15 minutes. The resulting pellet was washed with 75% ethanol, resuspended in 200 μl of RNase-free water and stored at −70°C.

#### Larvae

Each pooled sample of 60 larvae was macerated directly in the collection tube with disposable pestles. RNA was extracted with the High Pure Kit Nucleic Acid Kit DNA-RNA (Roche Diagnostics) according to the manufacturer’s recommendations. To determine the integrity and quality of extracted RNA, aliquots of randomly selected samples were quantified by fluorometry (Qubit 2.0 Invitrogen) and analyzed by electrophoresis on denaturing agarose gels. For each larval and adult sample, cDNA was synthesized using 500 μg of total RNA and reagents from the Transcriptor First Strand cDNA Synthesis kit (Roche Diagnostics), using the following thermal profile: 10 min., 25°C; 30 min., 55°C; 5 min., 85°C.

### Internal amplification control

Before performing the viral diagnosis, to ensure that lack of amplification was not due to poor extraction or the presence of PCR inhibitors, the samples were subjected to amplification of a 184 bp fragment of *Apis mellifera* Beta Actin gene, according to the protocol reported by Chen et al., (2005) [48], but adapted to a quantitative PCR protocol with SYBR green. The size of the amplified fragment was verified by electrophoresis on 2% agarose gel, in TAE1X, visualized under UV light on a transilluminator (NyxTechnik) and compared with GeneRuler ladder of 100-1000 bp (Thermo Scientific).

### Detection of viral pathogens by real-time PCR with SYBRGreen

We followed protocols reported by other authors for the detection of the targeted viruses [48–54] (Table S1). In this study, these protocols were adjusted to a real-time format with Green Essential FastStart Master kit (Roche Diagnostics), in a Nano LightCycler (Roche Diagnostics). Data were analyzed in the LightCycler SW 1.0 software. The results were interpreted as absence or presence of the amplified product, without performing quantifications. PCR techniques were done in separate reactions for each virus (Single PCR) with an aliquot of the cDNA from each larval and adult bee samples. Positive controls consisted of PCR products cloned in plasmids, which were obtained from the USDA-ARS Bee Research Laboratory (Beltsville MD, USA) and the Entomology Department, Volcani Center (Bet Dagan, Israel). Negative controls consisted of ultrapure water. Positive viral detections were verified by temperature melting curves analysis. Amplified viral fragments had the following lengths and melting point temperatures: DWV 700 bp, 82°C [55]; ABPV, 452 bp, 80°C [49]; SBV 340 bp, 83.7 [51], BQCV 284 bp, 80.5°C [50]. We followed two criteria to determine a sample as positive: a) confirmation of a single peak of the melting curve and b) the correct size of the amplified fragment. Additionally, amplicons of positive samples were sequenced and compared to NCBI database using Blast-n. The prevalence of virus was defined as the ratio between the number of PCR-positive to the total number of (colony-level) samples.

### Determining the prevalence of virus

The sampling was designed to be a broad survey not only searching for specific disease symptoms but also the presence and absence of the virus. Therefore, the virus-positive samples obtained by RT-PCR were interpreted as infected, but it does not imply that the originating material showed bees with overt illness. The prevalence of each virus was defined as the ratio between: Number of infected colonies / total number of sampled hives. For each virus, the prevalence was established in each region and the subsequent mean for all the three regions. We followed two criteria to determine a sample as positive: a) confirmation of a single peak of the melting curve and b) the correct size of the amplified fragment. Additionally, amplicons of positive samples were sequenced and compared to NCBI database using Blast-n.

### Statistical analysis

Initial analyses of normality and homoscedasticity of the VIL and viral prevalence values was conducted using the Kolmogorov-Smirnov and Levene tests [56]. Considering that the distributions of these values did not fulfill the assumption of parametric statistics, they were analyzed using the Kruskal Wallis and Mann Whitney U test [57]. Correlations among viral prevalence, altitude and season were conducted using Spearman’s nonparametric correlation [57, 58] and these values were used to construct a dissimilarity matrix (dissimilarity = 1 - correlation coefficient). The analyses of VIL and prevalence of each virus were calculated per region using the colony as a unit. Because Magdalena was sampled only during the rainy season and Sucre only during the dry season, we tested effects of season within the Boyacá region only. Similarly, we analyzed the effects of altitude within each region to avoid confounding the effects of altitude with those of season.

## Results

### Prevalence and infestation level of *Varroa destructor*

Of the 483 sampled colonies, finding that the vast majority were infested with *V. destructor* mites. Prevalence of *V. destructor* by region was as follows: Boyacá 89%, Magdalena 96% and Sucre 90%. *Varroa* infestation level (VIL) in Magdalena was higher than in Boyacá and Sucre (p= <0.0001), but levels did not differ significantly between Boyacá and Sucre (p=0.53, Figure 2).

**Figure 2.**
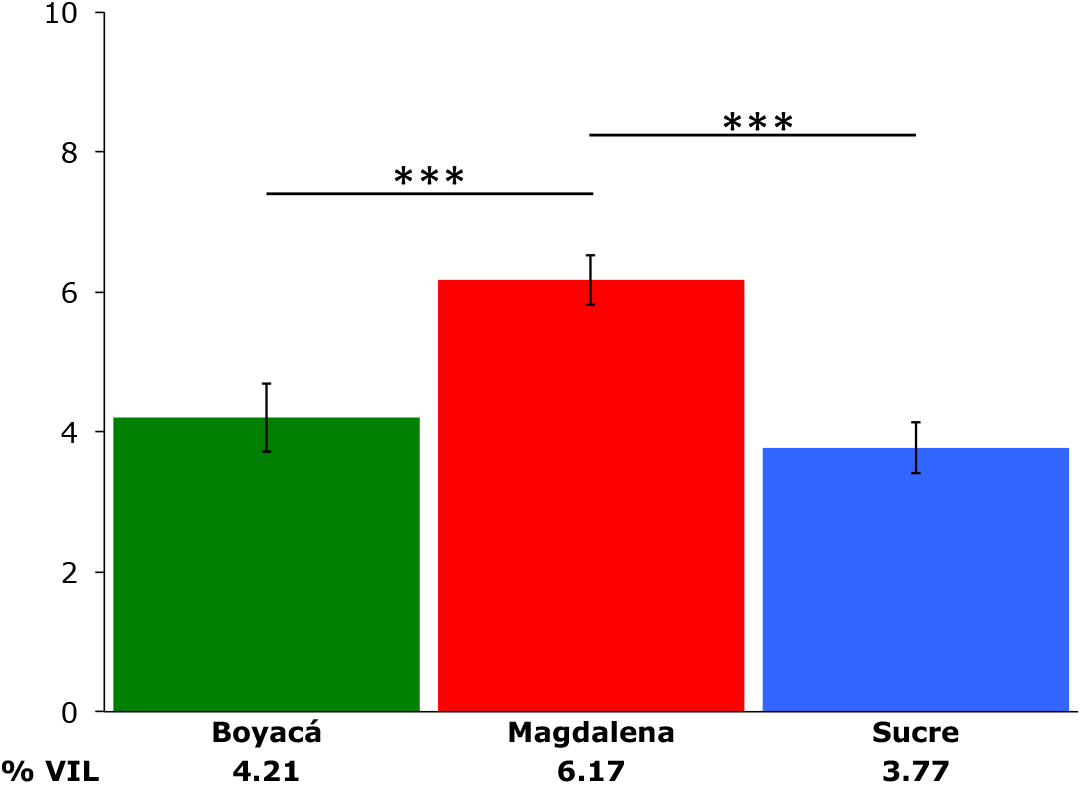
Varroa infestation levels per region. The y-axis indicates mean percentage of Varroa infestation. Error bars represent SE. Non-parametric data were analyzed with the Kruskal Wallis and Mann Whitney U test.

Overall, 68% of colonies showed values below 5%, 20.6% had VIL between 5 to 10%, 9% of colonies presented VIL between 10 to 15% and only 2.4% registered infestations level above of 15%. The region with the highest proportion of colonies with VIL above 5% was Magdalena. In contrast, Boyacá had a higher proportion of samples with VIL below 5% (Figure 3 and Table S2). We found significantly higher VIL (Mann-Whitney U test p<0.001) in the samples collected during the rainy season (mean 5.0, SE 0.312) compared with the samples collected during the dry season (mean 4. 1, SE 0.271). However, the effects of season are confounded with those of region and altitude, given that all samples from the relatively high-altitude Magdalena region were collected during the rainy season, whereas all samples from the low-altitude Sucre region were collected during the dry season.

**Figure 3.**
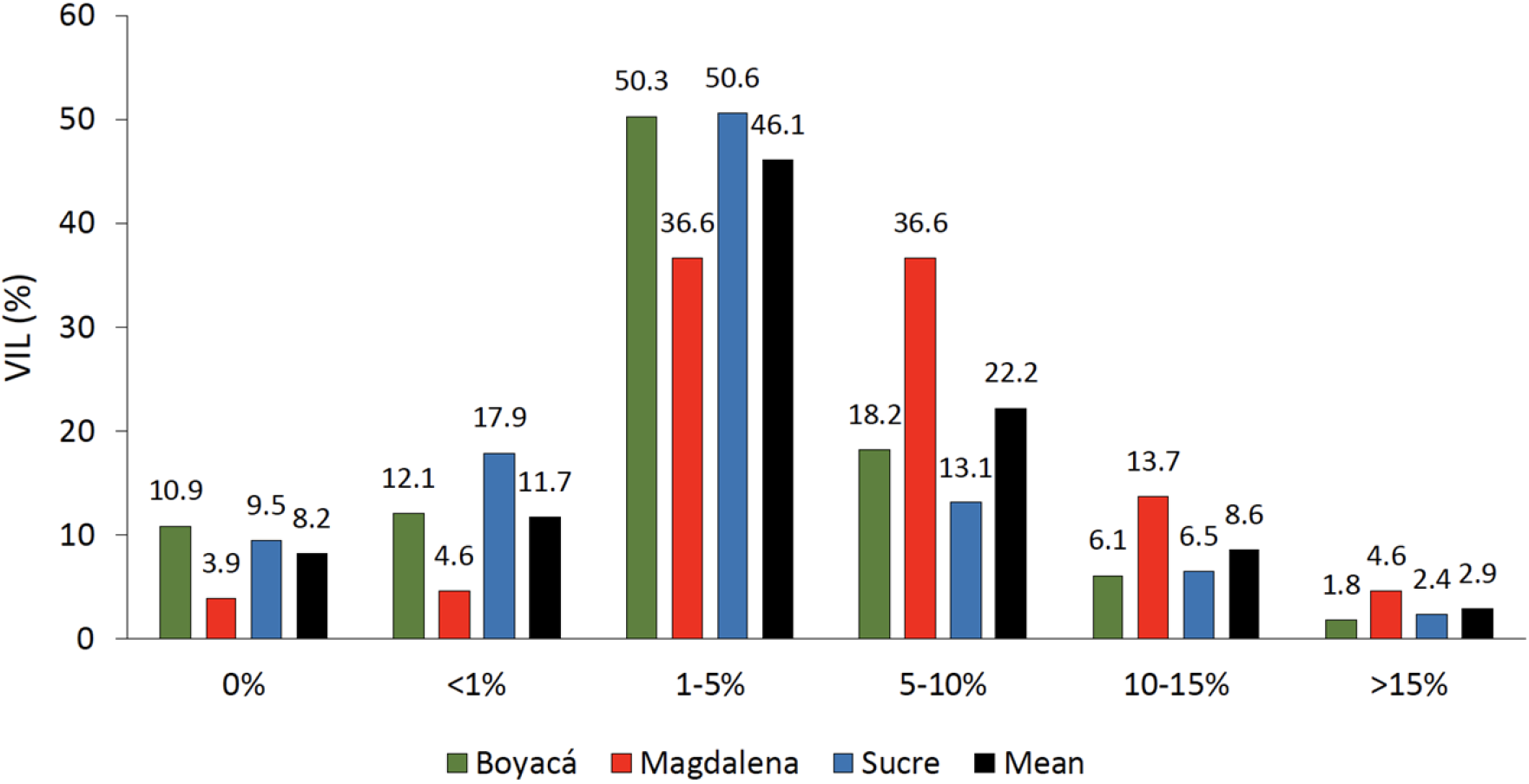
Histogram depicting percentages of samples with different percentages of VIL (mites per 100 adult bees).

### Viral co-infection

Of the seven viral pathogens tested, four of them (DWV, BQCV, SBV, ABPV) were found in both samples of larvae and adults from all three regions. The other three viruses tested (CBPV, IAPV, KBV) were not detected in any of the samples. Of the 483 colonies analyzed, we found no viruses in 35%, one virus in 31%, two viruses in 22.8%, three viruses in 10.1%, and all four viruses in only 1.1% of the colonies (Table 1).

**Table 1.** Percentages of viral co-infection. Percentage of samples no infected, infected with a single virus or coinfected with two, three or four of the viruses detected (DWV, SBV, BQCV and ABPV).

| Detection | % | Number of samples |
| --- | --- | --- |
| No virus | 35 | 172 |
| 1 virus | 31 | 152 |
| 2 viruses | 22.8 | 112 |
| 3 viruses | 10.1 | 50 |
| 4 viruses | 1.1 | 5 |

### Viral prevalence

In adults, SBV showed the higher percentage of prevalence (24.03%), followed by DWV (19.76%), BQCV (17.31%) and ABPV (11%). In larvae, BQCV was the virus with the higher percentage of prevalence (21.18%), followed by SBV (18.13%), DWV (15.27%) and ABPV (3.87%) (Figure 4, Table S3). Comparison between adult and larvae, revealed similar trends in the cases of DWV and BQCV, where no significant differences in the prevalence of these viruses were observed between the two life stages. In contrast, significant differences were found between larvae and adult stages for ABPV (p<0.001) and SBV (p=0.023). In adults, significant differences among viral prevalence were observed among ABPV with BQCV (p=0.005), DWV and SBV (p<0.001), as well as between BQCV and SBV (p=0.009). In larvae, significant differences among viral prevalence were observed among ABPV with BQCV (p<0.001), DWV and SBV (p<0.001), and between BQCV with DWV (p<0.001) (Figure 4).

**Figure 4.**
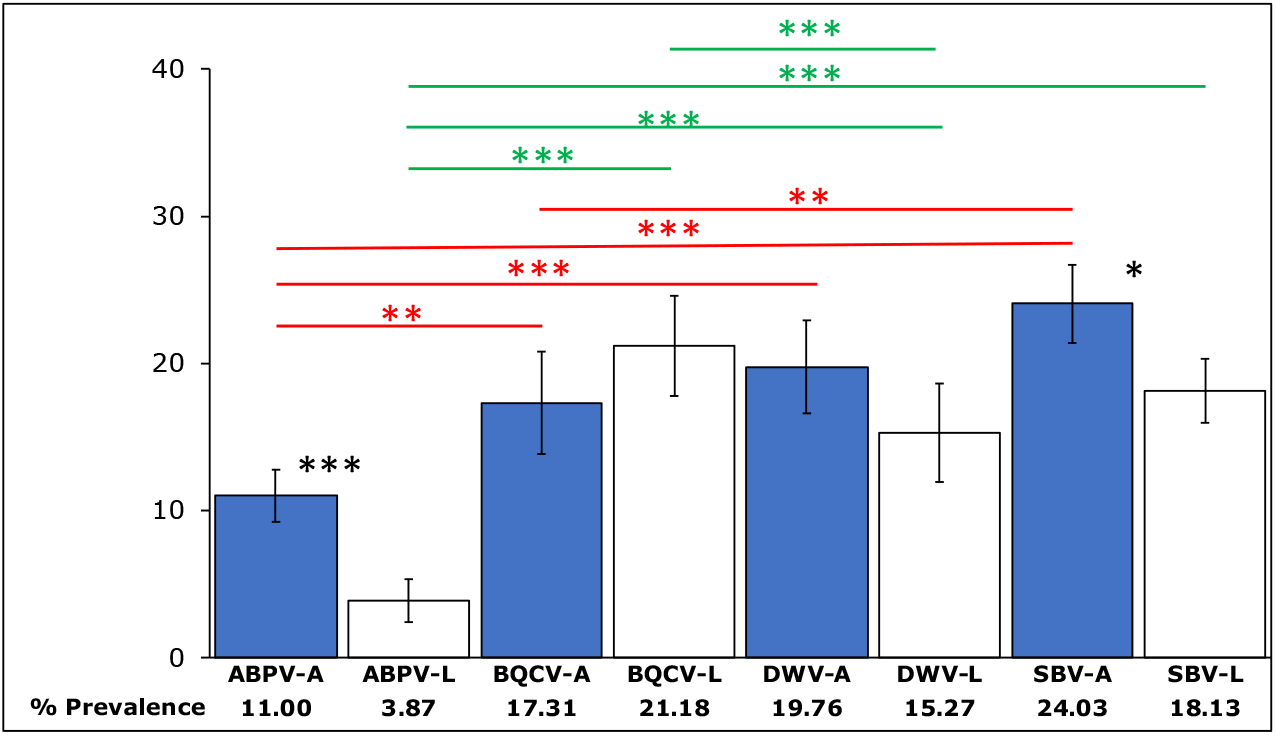
Overall percentages of viral prevalence (y-axis) in larvae (empty bars) and adult bees (filled blue bars). Significant differences in viral prevalence between larvae and adult per each virus and region are represented with black stars. Significant differences in the prevalence of the different viruses across regions in adults and larvae are represented by red and green starts, respectively (p< 0.05 *, p<0.001 **, p<0.0001***). Error bars represent SE.

Differences among regions were most evident in adults. Samples from Magdalena had the highest prevalence of the three most frequently detected viruses, whereas samples from Sucre had the lowest levels (Figure 5). Significant differences were found in the prevalence of the following viruses between the following regions: Boyacá and Magdalena: ABPV (p<0.001), BQCV (p=0.04) and SBV (p<0.001). Magdalena and Sucre: ABPV (p<0.001), BQCV (p<0.001), DWV (p=0.002) and SBV (p<0.001). Boyacá and Sucre: BQCV, DWV and SBV (p<0.001) (Figure 5). Among larvae, a similar pattern than the observed in adults was found for SBV. However, prevalence of larval DWV was highest in Boyacá rather than in Magdalena, but likewise low in Sucre (Figure 6). Significant differences in larval viral prevalence between regions were as follows: Boyacá and Magdalena: DWV (p=0.01) and SBV (p<0.001). Magdalena and Sucre: DWV (p=0.024) and SBV (p<0.001). Boyacá and Sucre: DWV (p<0.001) (Figure 6).

**Figure 5.**
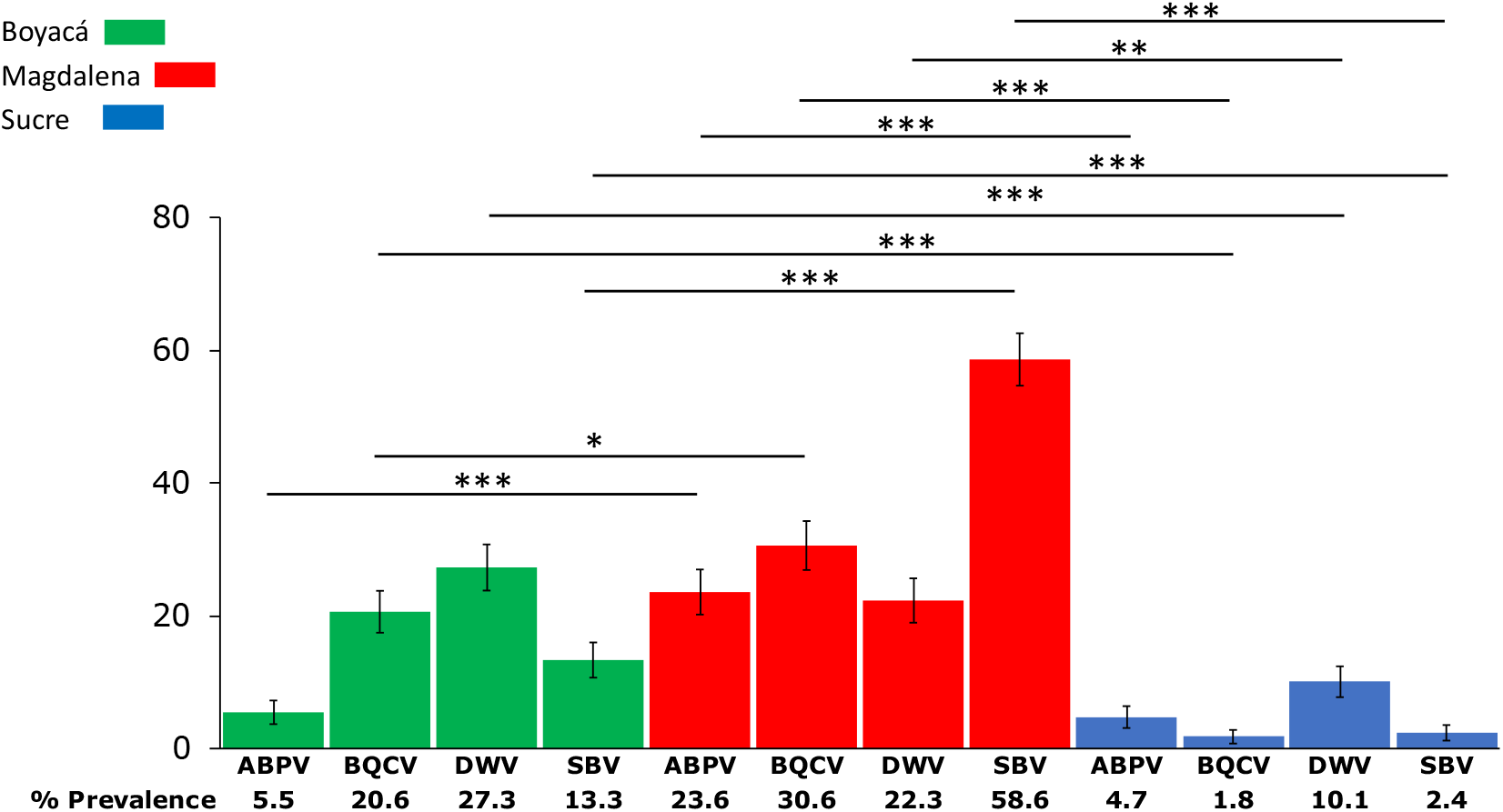
Percentages of viral prevalence (y-axis) and adult bees in Boyacá (red bars), Magdalena (green bars) and Sucre (blue bars). Significant differences in viral prevalence per each virus across regions are indicated with black stars (p< 0.05 *, p<0.001 **, p<0.0001***). Error bars represent SE.

**Figure 6.**
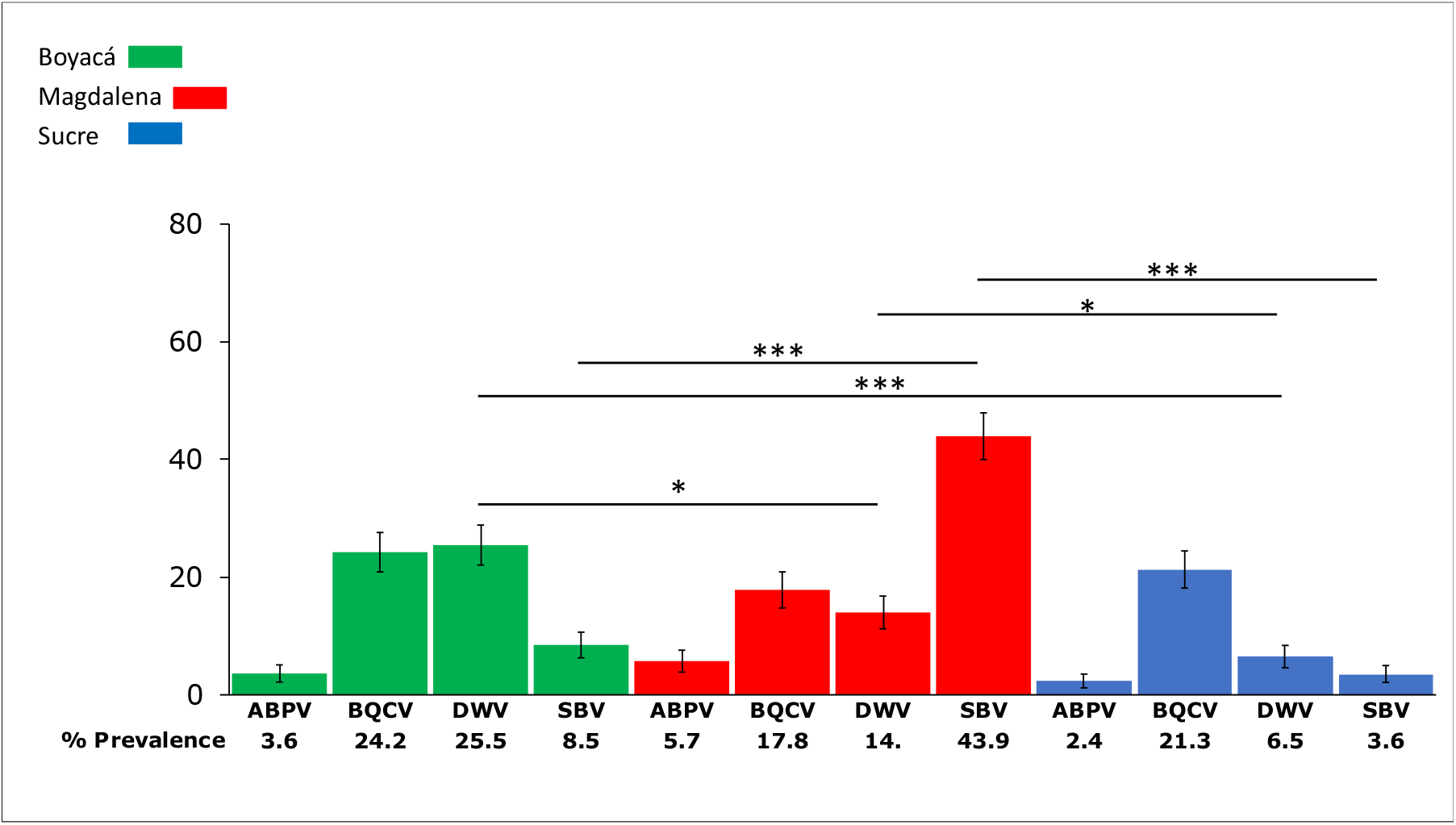
Percentages of viral prevalence and larvae in Boyacá (red bars), Magdalena (green bars) and Sucre (blue bars). Significant differences in viral prevalence per each virus across regions are indicated with black stars (p< 0.05 *, p<0.001 **, p<0.0001***). Error bars represent SE. Data was analyzed using one-way ANOVA followed by Bonferroni correction.

### Correlations among variables

#### Correlations among viral prevalence

We first determined the associations concerning the prevalence of the different viruses, pooled across the three regions. Overall analysis of viral prevalence between larval and adult stages for each virus, showed positive correlations for all the viruses analyzed with the exception of BQCV. In adults, we found significant positive correlations between each of the four viruses. In larvae, significant positive correlations were observed between ABPV and DWV, ABPV and SBV and BQCV and SBV (Table 2).

**Table 2.**
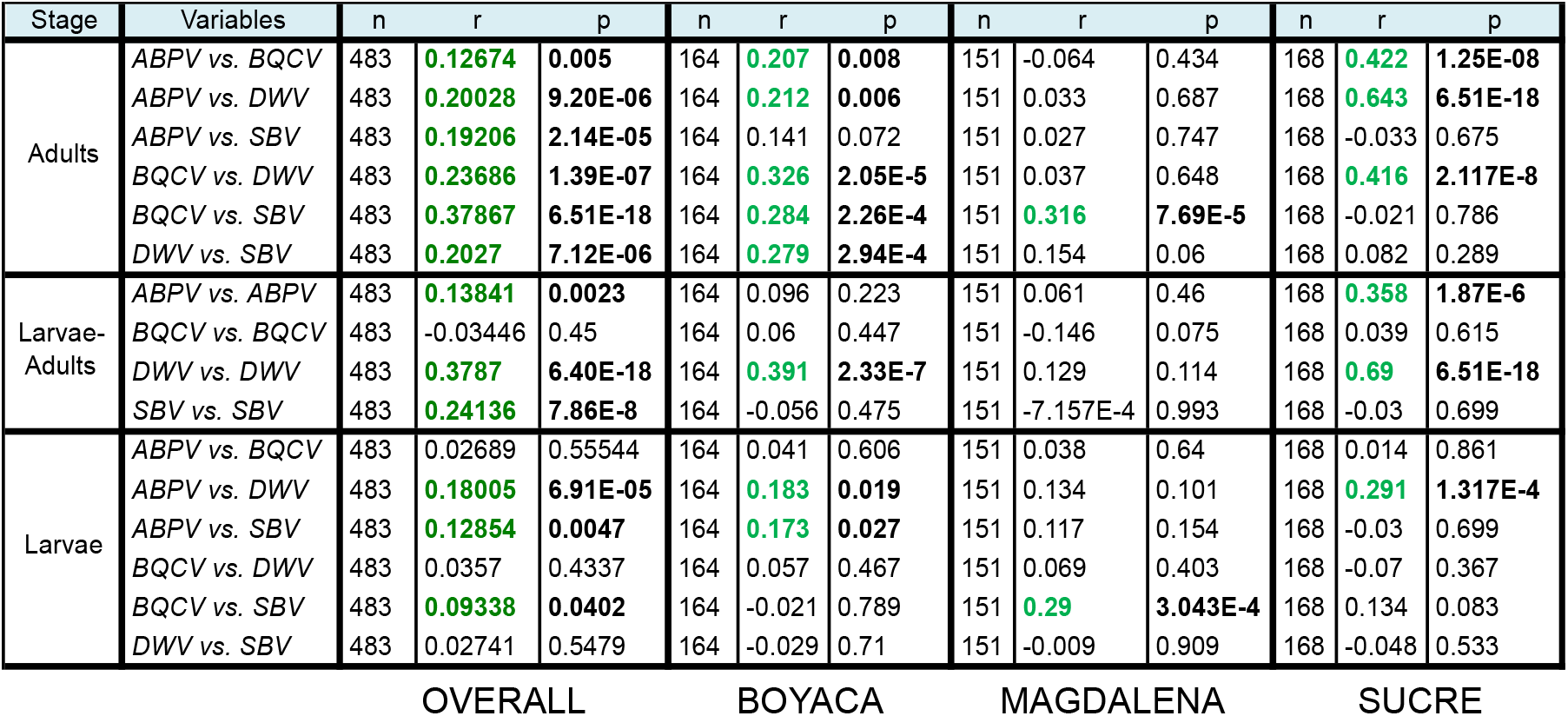
Wise Spearman rank correlations among viral prevalence in adults and larvae. Significant differences (p<0.05) are represented in bold case. Positive correlations are showed in green case.

| Stage | Variables | n | r | p | n | r | p | n | r | p | n | r | p |
| --- | --- | --- | --- | --- | --- | --- | --- | --- | --- | --- | --- | --- | --- |
| Adults | ABPV vs. BQCV | 483 | **0.12674** | **0.005** | 164 | **0.207** | **0.008** | 151 | -0.064 | 0.434 | 168 | **0.422** | **1.25E-08** |
|  | ABPV vs. DWV | 483 | **0.20028** | **9.20E-06** | 164 | **0.212** | **0.006** | 151 | 0.033 | 0.687 | 168 | **0.643** | **6.51E-18** |
|  | ABPV vs. SBV | 483 | **0.19206** | **2.14E-05** | 164 | 0.141 | 0.072 | 151 | 0.027 | 0.747 | 168 | -0.033 | 0.675 |
|  | BQCV vs. DWV | 483 | **0.23686** | **1.39E-07** | 164 | **0.326** | **2.05E-5** | 151 | 0.037 | 0.648 | 168 | **0.416** | **2.117E-8** |
|  | BQCV vs. SBV | 483 | **0.37867** | **6.51E-18** | 164 | **0.284** | **2.26E-4** | 151 | **0.316** | **7.69E-5** | 168 | -0.021 | 0.786 |
|  | DWV vs. SBV | 483 | **0.2027** | **7.12E-06** | 164 | **0.279** | **2.94E-4** | 151 | 0.154 | 0.06 | 168 | 0.082 | 0.289 |
| Larvae-Adults | ABPV vs. ABPV | 483 | **0.13841** | **0.0023** | 164 | 0.096 | 0.223 | 151 | 0.061 | 0.46 | 168 | **0.358** | **1.87E-6** |
|  | BQCV vs. BQCV | 483 | -0.03446 | 0.45 | 164 | 0.06 | 0.447 | 151 | -0.146 | 0.075 | 168 | 0.039 | 0.615 |
|  | DWV vs. DWV | 483 | **0.3787** | **6.40E-18** | 164 | **0.391** | **2.33E-7** | 151 | 0.129 | 0.114 | 168 | **0.69** | **6.51E-18** |
|  | SBV vs. SBV | 483 | **0.24136** | **7.86E-8** | 164 | -0.056 | 0.475 | 151 | -7.157E-4 | 0.993 | 168 | -0.03 | 0.699 |
| Larvae | ABPV vs. BQCV | 483 | 0.02689 | 0.55544 | 164 | 0.041 | 0.606 | 151 | 0.038 | 0.64 | 168 | 0.014 | 0.861 |
|  | ABPV vs. DWV | 483 | **0.18005** | **6.91E-05** | 164 | **0.183** | **0.019** | 151 | 0.134 | 0.101 | 168 | **0.291** | **1.317E-4** |
|  | ABPV vs. SBV | 483 | **0.12854** | **0.0047** | 164 | **0.173** | **0.027** | 151 | 0.117 | 0.154 | 168 | -0.03 | 0.699 |
|  | BQCV vs. DWV | 483 | 0.0357 | 0.4337 | 164 | 0.057 | 0.467 | 151 | 0.069 | 0.403 | 168 | -0.07 | 0.367 |
|  | BQCV vs. SBV | 483 | **0.09338** | **0.0402** | 164 | -0.021 | 0.789 | 151 | **0.29** | **3.043E-4** | 168 | 0.134 | 0.083 |
|  | DWV vs. SBV | 483 | 0.02741 | 0.5479 | 164 | -0.029 | 0.71 | 151 | -0.009 | 0.909 | 168 | -0.048 | 0.533 |

OVERALL
BOYACA
MAGDALENA
SUCRE

The extent of covariation in viral prevalence varied across regions and life stages. In adults from Boyacá, all comparisons among viruses were significant with the exception of ABPV and SBV. However, in Magdalena, only BQCV and SBV were significantly correlated. In within-region analyses of larvae, correlations among viruses were less pronounced than in adults. Two of six pairs of viruses were correlated in Boyacá, and one of six pairs each in Magdalena and Sucre (Table 2).

**Figure 7.**
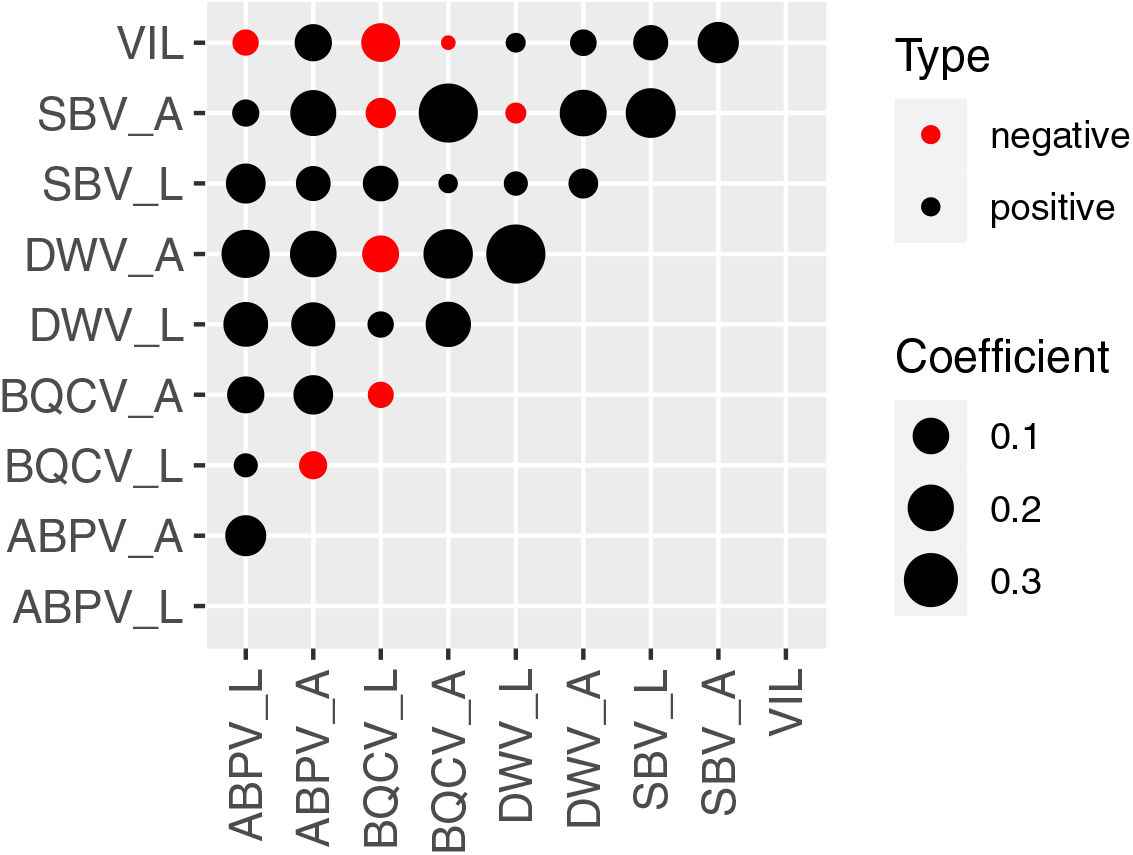
Correlations among varroa infestation levels and viral prevalence in adults (A) and larvae (L). The circle size is proportional to the correlation coefficient. Blue circles indicate positive correlations; red circles indicate negative correlations.

#### Varroa infestation levels

In adults of all regions combined, we found significant positive correlations between VIL and prevalence of ABPV (r=0.106, p<0.05) and SBV (r=0. 144, p<0.05). On the other hand, VIL did not correlate significantly with DWV or BQCV (Table3). In larvae, the only correlation was a negative association VIL and BQCV (r= −0.115, p<0.05). Within each region, significant correlation was observed between VIL and viral prevalence only in larvae from Boyacá, where VIL was negatively correlated with BQCV (r= −0.226, p<0.01); no within-region correlations were significant in adults (Table 3).

**Table 3.**
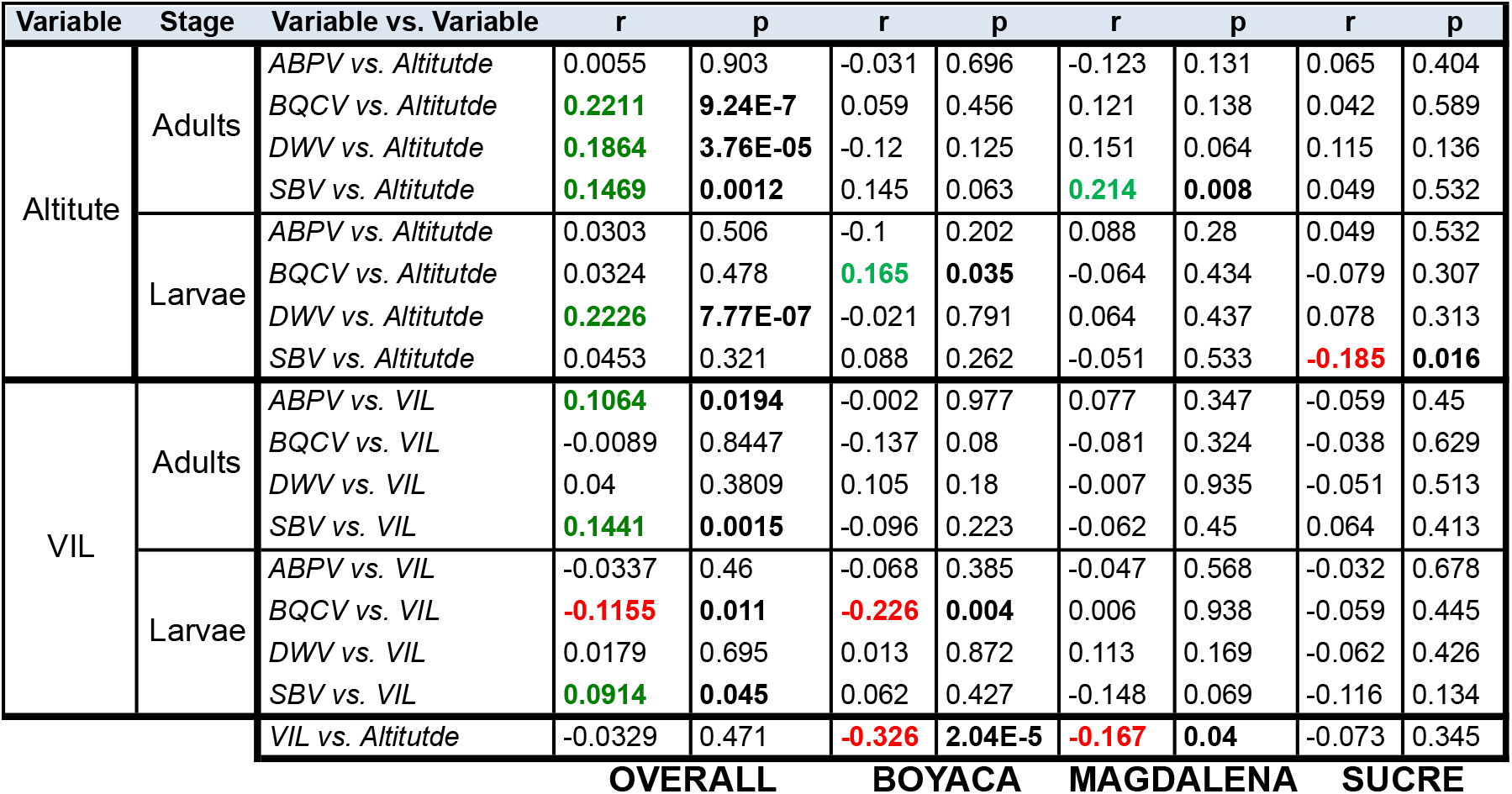
Wise Spearman rank correlations among VIL, altitude and viral prevalence in adults and larvae. Significant differences (p<0.05) are represented in bold case. Positive and negative correlations are showed in green and red case, respectively.

#### Altitude

We found no correlation between altitude and VIL across the three regions combined. However, in within-region analyses that removed the confounding effect of season, altitude was negatively correlated with VIL within both Boyacá (r= −0.326, p<0.001) and Magdalena (r= −0.167, p<0.05) (Table 3). In contrast, we found positive correlations between altitude and viral prevalence for three of four viruses in adults (BQCV (r= 0.221, p<0.001) DWV (r= 0.186, p<0.001), and SBV (r= 0.146, p<0.01), and for DWV in larvae (r= 0.222, p<0.001) (Table 3).

#### Season

Seasonal analysis was restricted to the Boyacá, the only region where samples were collected both during the raining and dry season. Our results showed significant positive correlations between the rainy season and the prevalence of BQCV, ABPV, while DWV showed a positive correlation in the limit of statistical significance (r=0.153, p=0.05). On the other hand, in larvae, only DWV and BQCV exhibited significant positive correlation with the rainy season.

**Table 4.** Wise Spearman rank correlations among viral prevalence and rainy season in adults and larvae. Significant differences (p<0.05) are represented in bold case. Positive and negative correlations are showed in green and red case, respectively

| Age | Variable vs. Variable | n | r | p |
| --- | --- | --- | --- | --- |
| Adults | ABPV_A vs. Rains | 164 | 0.253 | **0.001** |
|  | BQCV_A vs. Rains | 164 | 0.266 | **5.78E-4** |
|  | DWV_A vs. Rains | 164 | 0.153 | **0.05** |
|  | SBV_A vs. Rains | 164 | -0.052 | 0.505 |
| Larvae | ABPV_L vs. Rains | 164 | 0.075 | 0.343 |
|  | BQCV_L vs. Rains | 164 | 0.426 | **1.32E-8** |
|  | DWV_L vs. Rains | 164 | 0.28 | **2.76E-4** |
|  | SBV_L vs. Rains | 164 | -0.204 | **0.009** |

## Discussion

### *Varroa* prevalence and infestation levels

Our results revealed a high *V. destructor* prevalence (92%). Dissemination of *V. destructor* has increased considerably in South America since the 1990s. This phenomenon is exemplified by the case of EHB populations in Chile, where *Varroa* prevalence increased from 80% in 2007 [59] to 93% in 2013 [46]. High percentages of varroa prevalence also have been reported in AHB populations in southern Brazil (95.7%) [60], which are consistent with the prevalence observed in the present study (92%). However, *Varroa* prevalence levels have been maintained at lower levels in other countries with either predominant AHB or EHB populations, such as Uruguay (75.7%) [38] and Argentina (74%) [46], respectively. Thus, at present, there is not an evident trend between the degree of Africanization and prevalence of *Varroa*, suggesting that other factors, including management, may modulate observed prevalence levels.

Although prevalence was high, *Varroa* infestation level (VIL) was relatively low (4.6%). There have been few previous studies on VIL in Latin American countries. These include countries with EHB populations, such as Chile, with VIL of 5-9% [46, 61], and countries with predominant AHB populations (~80% African mitotypes) such as Uruguay [62] and Mexico [63], with VIL of 7.5% [64] and 5-7.4% [65, 66], respectively. Relative to these regional neighbors, Colombia appears to have the lowest VIL (4.6%), despite being the only country from this group where acaricides are not routinely used. Among these countries, Colombia also has the highest percentage of AHB (98.3%) [44]. suggesting a negative relationship between VIL and the proportion of AHBs in the population. Further studies in additional regions with comparable climates are required to confirm the generality of this trend.

For comparison, in the United States, *V. destructor* prevalence and infestation levels for colonies used in stationary beekeeping operations during 2009-2014 was 97.0% and 5.99% [13]. Although these values are only marginally higher compared with what we reported for Colombia, a critical difference is that AHBs in our study did not receive acaricide treatments. [67] Furthermore, it is also interesting to note that while in the United States and Canada [7, 13] *V. destructor* is regarded as a major contributor to yearly losses that have exceeded 40% annually in the United States from 2015-2016 [68]; in Colombia, varroosis – the disease caused by Varroa infestation - is not considered an important problem for beekeepers and yearly colony losses were estimated at 10.8% [69].

Previous studies comparing VIL between regions with tropical and temperate climates in Mexico have found either higher VIL in tropical regions [66], or no significant differences between climates [65]. In our study, Boyacá is the region with the higher altitude, followed by Magdalena and Sucre. Thus, Boyacá has a relatively colder climate, Magdalena region, has an intermediate subtropical temperature, and Sucre a tropical climate. In our study, overall analysis of the three regions, do not show significant correlations between VIL and altitude. However, regional analysis to remove the confounding effect of region and season revealed a significant negative correlation between VIL and altitude in the two regions with both higher altitude and larger variation in the altitude (Boyacá and Magdalena). In contrast, no significant correlation between VIL and altitude was found in the region with lower altitude and smaller difference in this variable (Sucre). Altogether, these results support the notion that altitude is an important factor influencing negatively VIL in tropical and neotropical regions. Thus, this effect of altitude on Varroa infestation resembles the reduction on Varroa population observed in honey bee colonies before the winter in non-temperate geographic regions.

### Viral prevalence

In colonies from the three surveyed regions, the presence of four viral pathogens was detected. However, prevalence was lower than that reported for other countries in South America [20, 46] and Mexico [66], and none of the colonies in this study showed evident symptoms of overt infection, suggesting that infection intensity was low. The Magdalena region had the highest VIL as well as the highest prevalence of ABPV, BQCV and SBV. In contrast, Sucre had the lowest VIL and prevalence of the four viruses. These results suggest a possible association between *V. destructor* infestation and viral prevalence. Our analysis shows that there is a positive correlation between VIL and the prevalence of ABPV and SBV. However, we found no significant correlations between the prevalence of VIL with DWV or with BQCV. Although this last result was unexpected, it is interesting to note that the bees in this study did not show evident symptoms of overt infection. Studies showing positive association between DWV and VIL were conducted in colonies with higher levels of *Varroa* infestation or in parasitized individuals [10, 66]. It remains to be further investigated if this lack of association is related to the low VIL observed in Colombian AHBs.

The global spread of *V. destructor* has selected for and disseminated highly infectious and pathogenic DWV strains in European-derived populations around the world [70].. In the United States and Europe, pathogenic viruses have increased their prevalence and virulence, changing from asymptomatic to evident symptoms of infection. Comparing viral infection levels in the United States with those we are reporting as present in Colombia illustrates a sharp contrast. The reported prevalence of three viral pathogens screened in the United States during 2009-2014 was: DWV 85.09 %, ABPV 21.74 %, and BQCV 90.03 % [13]. Interestingly, although both Colombia and the United States shared a similar VIL, the prevalence of surveyed viral pathogens in Colombia is considerably lower.

We found highly significant positive correlation between altitude and the prevalence of DWV, BQCV and SBV in adult bees. Our results are consistent with the finding of higher DWV prevalence in colonies from temperate climate compared with those from tropical regions in Mexico [66]. These results support the proposal that cold stress could weaken the immune response and enhance viral replication. Supporting this possibility, cold temperatures have been associated with down-regulation of the cellular immune response [71] and higher DWV titers [10].

Seasonal differences are an important factor to be considered in the study of VIL and viral prevalence. In countries with temperate climates, higher VIL and viral titers are found during the fall [13, 20]. Regions near the equator do not experience substantial fluctuations in temperature. However, these tropical regions experienced important seasonal differences in pluvial precipitation and two distinctive dry and rainy season can be distinguished during the year. One important caveat of the present study is that our experimental design was not specifically planned to uncouple the effect of pluvial precipitation during the year. Despite this limitation, samples were collected during both the dry and raining seasons in Boyacá, the region with the higher difference in altitude. This allowed for an initial estimation of the effect of pluvial precipitations on VIL and viral prevalence as well as the interactions among these variables with altitude. Correlation analysis showed a significant positive association between the raining season and viral prevalence. These results suggest that pluvial precipitations have an important effect on viral prevalence in AHB populations of tropical regions.

### Conclusive remarks

In this report, we documented low *Varroa* infestation levels in non-acaricide treated, fully Africanized Colombian honey bee populations. These results are consistent with previous studies that found low *Varroa* infestation levels in Africanized populations of other Latin American countries, and lesser infestation levels with increasing proportions of Africanized relative to European bees [46]. Moreover, despite lack of treatment with acaricides, Colombian bees had low prevalence and exhibited no obvious symptoms of the most pervasive honey bee viruses. Further studies are required for a better understanding of the interactions between *Varroa* and virus in AHBs, including determination of viral titers and the effect of season. This study provides valuable insights into understanding the relationship between *V. destructor* and the transmission and spread of honey bee viruses in AHBs.

## Acknowledgments

The authors would like to thank Edgar Arias, Carlos Baez, Viviana Gamboa, Umberto Moreno and Rogelio Rodríguez for assistance in the collection and processing of the samples. We are grateful with the Conservationist Beekeepers Association of the Sierra Nevada de Santa Marta, the Rural Beekeepers Association of Sucre and the Beekeepers Association of Boyacá for their collaboration during sampling. We are also grateful to Jay Evans, Yanping Chen and Nor Chejanovsky for scientific advice. We would also like to acknowledge Belen Branchiccela, Eugene Ryabov and Mohamed Alburaki for reviewing the manuscript.

## Author Contributions

Conceived and designed the experiments: VMT and JF. Performed the experiments: VMT and AS. Analyzed the data: MC, VMT, EP-Y, VMS and HS. Contributed reagents/materials/analysis tools: JF and FA. Wrote the paper: MC, VMT, EP-Y and SM.

